# Polymorphism-aware species trees with advanced mutation models, bootstrap and rate heterogeneity

**DOI:** 10.1101/483479

**Authors:** Dominik Schrempf, Bui Quang Minh, Arndt von Haeseler, Carolin Kosiol

**Affiliations:** Department of Biological Physics, Eötvös Loránd University, Budapest, Hungary; Centre for Biological Diversity, University of St Andrews, United Kingdom; Ecology and Evolution, Research School of Biology, Australian National University, Australia; Center for Integrative Bioinformatics Vienna, Max F. Perutz Laboratories, University of Vienna, Medical University of Vienna, Austria; Bioinformatics and Computational Biology, Faculty of Computer Science, University of Vienna, Austria; Institut für Populationsgenetik, Vetmeduni Vienna, Austria

**Keywords:** Incomplete Lineage Sorting, Species Tree, Phylogenetics, Polymorphismaware Phylogenetic Model, Boundary Mutation Model

## Abstract

Molecular phylogenetics has neglected polymorphisms within present and ancestral populations for a long time. Recently, multispecies coalescent based methods have increased in popularity, however, their application is limited to a small number of species and individuals. We introduced a polymorphism-aware phylogenetic model (PoMo), which overcomes this limitation and scales well with the increasing amount of sequence data while accounting for present and ancestral polymorphisms. PoMo circumvents handling of gene trees and directly infers species trees from allele frequency data. Here, we extend the PoMo implementation in IQ-TREE and integrate search for the statistically best-fit mutation model, the ability to infer mutation rate variation across sites, and assessment of branch support values. We exemplify an analysis of a hundred species with ten haploid individuals each, showing that PoMo can perform inference on large data sets. While PoMo is more accurate than standard substitution models applied to concatenated alignments, it is almost as fast. We also provide `bmm-simulate`, a software package that allows simulation of sequences evolving under PoMo. The new options consolidate the value of PoMo for phylogenetic analyses with population data.

## 1 Introduction

Molecular phylogenetics seeks to estimate evolutionary relationships of species depicted as species trees by modeling the change and development of hereditary sequences. Established methods (e.g., Tavaré, 1986; Yang, 2006) that neglect molecular variation on the population level suffer from incongruencies between genomic regions which may arise due to incomplete lineage sorting (ILS, Knowles, 2009; Maddison, 1997). The effect of ILS becomes large if the considered species are closely related or if internal branches of the species tree are short when measured in number of generations divided by the effective population size Pamilo and Nei (1988). Short branches, especially when appearing in caterpillar-like topologies may lead to statistical inconsistency Degnan (2013); Degnan and Rosenberg (2009) of approaches such as concatenation Gadagkar *et al.* (2005), where sequences for different genes of the same individual are joined to form one overall alignment.

Consequently, development of phylogenetic methods has increasingly focused on explicit modeling of population genetic effects Leaché and Oaks (2017). Most methods employ the multispecies coalescent model Rannala and Yang (2003) to reconcile gene trees, i.e., the evolutionary histories of genes, with the species tree. The species tree is either jointly estimated with the gene trees (e.g., Drummond and Rambaut, 2007; Heled and Drummond, 2010; Liu, 2008) or reconstructed from previously estimated gene trees (e.g., Liu *et al.*, 2010; Mirarab *et al.*, 2014).

We have recently proposed a polymorphism-aware phylogenetic model (PoMo, De Maio *et al.*, 2015; Schrempf *et al.*, 2016) for estimating species trees from genome-wide data for up to dozens of species with multiple individuals each, which bypasses the computational burden of estimating gene trees. PoMo can be viewed as an extension of classical substitution models which additionally considers polymorphisms. Present as well as ancestral polymorphisms are described along a species tree by separating mutation and genetic drift in a population genetic framework that we have termed multivariate boundary mutation model Schrempf and Hobolth (2017); Vogl and Clemente (2012). The substitution model (e.g., HKY model, Hasegawa *et al.*, 1985) extended by the multivariate boundary mutation model is referred to as mutation model, because frequency changes are considered separately. PoMo improves estimation of mutation parameters and species trees compared to established methods such as concatenation when ILS has blurred the phylogenetic signal. Extensive evaluations showed good performance of PoMo against other methods De Maio *et al.* (2015); Schrempf *et al.* (2016). Here we extend our implementation into the maximum likelihood software IQ-TREE Nguyen *et al.* (2015), referred to as IQ-TREE-PoMo, which includes several new features as detailed in the next section.

## 2 New Approaches

### Big data

In contrast to the multispecies coalescent model, PoMo scales much better with the number of analyzed species. The performance of IQ-TREE PoMo was tested by applying it to simulated multiple sequence alignments (MSA) of a length of up to 1 million nucleotides, and 100 species with ten sampled individuals each corresponding to gene trees with 1000 leaves (see Materials and Methods).

### Simulator

We developed and released a software package to generate sequences evolved under various boundary mutation models (bmms), in particular PoMos, called bmm-simulate (https://github.com/pomo-dev/bmm-simulate). Given a species tree, the simulator directly generates polymorphic sequences using the discrete multivariate boundary mutation model with mutation rate heterogeneity (see below). This simulation tool complements the coalescent simulators available which we have used in this and in previous studies and allows exact assessment of inference accuracy under PoMo.

### Advanced models

Heterogeneity of rates across sites is a consequence of varying mutation and fixation rates. If rate heterogeneity is not taken into account, sequence distance is underestimated and phylogenetic analyses may suffer from long branch attraction artifacts (Yang, 2006, p.19).

Therefore, we developed new theory for PoMo to incorporate Γ distributed mutation rate heterogeneity across sites Yang (1994). The new theory is necessary as only the mutation rates are heterogeneous across sites, while drift rates stay constant for our model (see Materials and Methods). A simulation study with bmm-simulate showed high accuracy when estimating the Γ shape parameter and other parameters on trees with twelve species.

Furthermore, species tree search can be performed with fixed parameters. Manual specification of parameters is useful when detailed knowledge is available or when likelihoods are compared between different analyses. In practice, we often do not have total control over the number of individuals in the population that are sampled and sequenced (e.g., sequences are retrieved from existing databases, re-sequencing runs failed or are simply too costly even for as few as 10 individuals). We therefore present two strategies, called weighted binomial and weighted hypergeometric sampling, to initialize the likelihoods of the PoMo states at the tips of the trees, and to account for this missing data problem.

### ModelFinder

Model selection, which includes finding the best-fit mutation model as well as a model of mutation rate heterogeneity across sites, is crucial when performing molecular phylogenetic analyses. The improved implementation of PoMo allows for highly flexible polymorphism-aware phylogenetic analyses in that the most suitable evolutionary model for the data at hand can be automatically determined using statistical model search with ModelFinder Kalyaanamoorthy *et al.* (2017). ModelFinder is a model selection framework in IQ-TREE which has now been made available for IQ-TREE-PoMo. The Bayesian Schwarz (1978) or the Akaike (1973) information criterion are employed to determine the best-fit model.

### Bootstrapping

IQ-TREE-PoMo is fast enough to allow for bootstrapping Efron (1979); Felsenstein (1985) on large data sets. Evaluation of branch support values using the branch-wise approximate likelihood ratio test (SH-aLRT, Guindon *et al.*, 2010) as well as standard nonparameteric bootstrap Efron (1979) and ultrafast bootstrap (UFBoot2, Hoang *et al.*, 2018) is now possible. The ability of bootstrapping was tested on a data set of great ape genomes (see Material and Methods, Prado-Martinez *et al.*, 2013).

## 3 Results

### Big data

The phylogenetic analysis of the simulated MSAs of 100 species with ten sampled individuals each shows that consideration of heterozygosity improves estimates considerably compared to the standard concatenation approach. Branch score distance (Fig. 1), as well as Robinson-Foulds distance (Fig. S1) strongly decrease, especially when sufficient data is available. Remarkably, IQ-TREE-PoMo already outperforms concatenation for as little as ten genes (10 000 sites) and exhibits branch score distances around ten times more accurate when 1000 genes (one million sites) are available. Further, the progression of branch score distance for the concatenation method with increasing amount of data is not monotonically decreasing. The average run time (wall-clock time) of IQ-TREE-PoMo for 1000 genes which corresponds to a sequence length of one million sites is 12.3 ± 0.7 hours on a 2.6 GHz processor with 16 physical cores (Intel^®^ Xeon^®^ CPU E5-2650 v2 @ 2.60GHz) The concatenation method is approximately six times faster (2 ± 0.25 hours).

**Figure 1:**
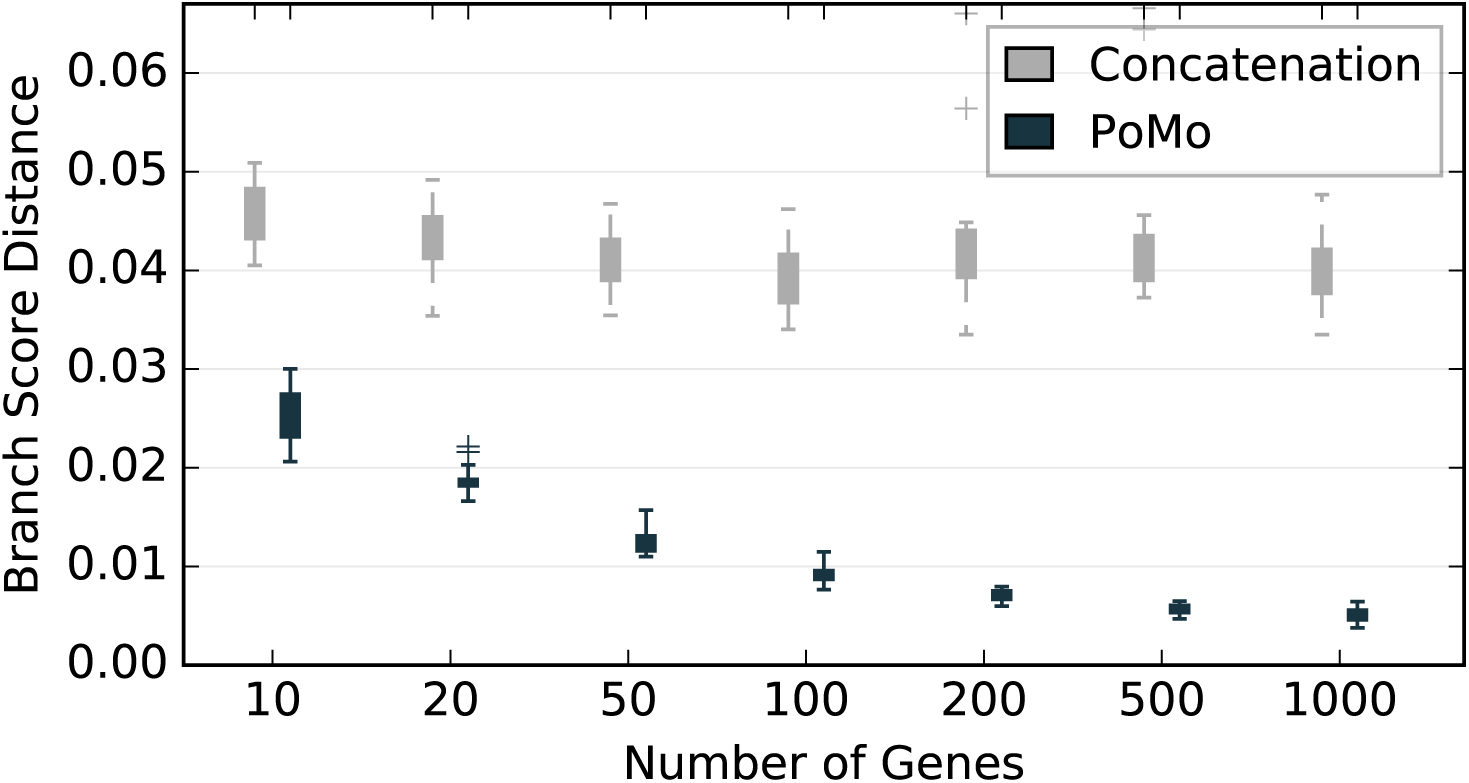
Branch score distance of concatenation approach and IQ-TREE-PoMo with *N* = 10 and weighted binomial sampling for Yule trees with 100 species and ten individuals each. The tree height measured in coalescent units is 6*N_e_*, where *N_e_* is the effective population size. The HKY model was used for both inference methods. The heterozygosity is *θ_W_* = 0.005 per site. Each gene spans 1000 sites. The error bars are standard deviations of ten replicate analyses.

### Advanced models

We assess the accuracy of estimating rate heterogeneity by focussing on the shape parameter α of the Γ distribution. The relative error (difference between true and estimated value in percent) of the shape parameter *α* for ten replicate analyses is usually within two percent (see Fig. 2 for α = 0.3, 1.0, and 5.0, respectively) but can be higher for more extreme *α* values (≈ 20 and (≈ 400 percent for *α* = 0.1, and 10, respectively; Fig. S2; – S4). The relative errors in terms of branch score distance and the transition to transversion ratio parameter of the HKY mutation model are mostly below 0.5 and one percent, respectively.

**Figure 2:**
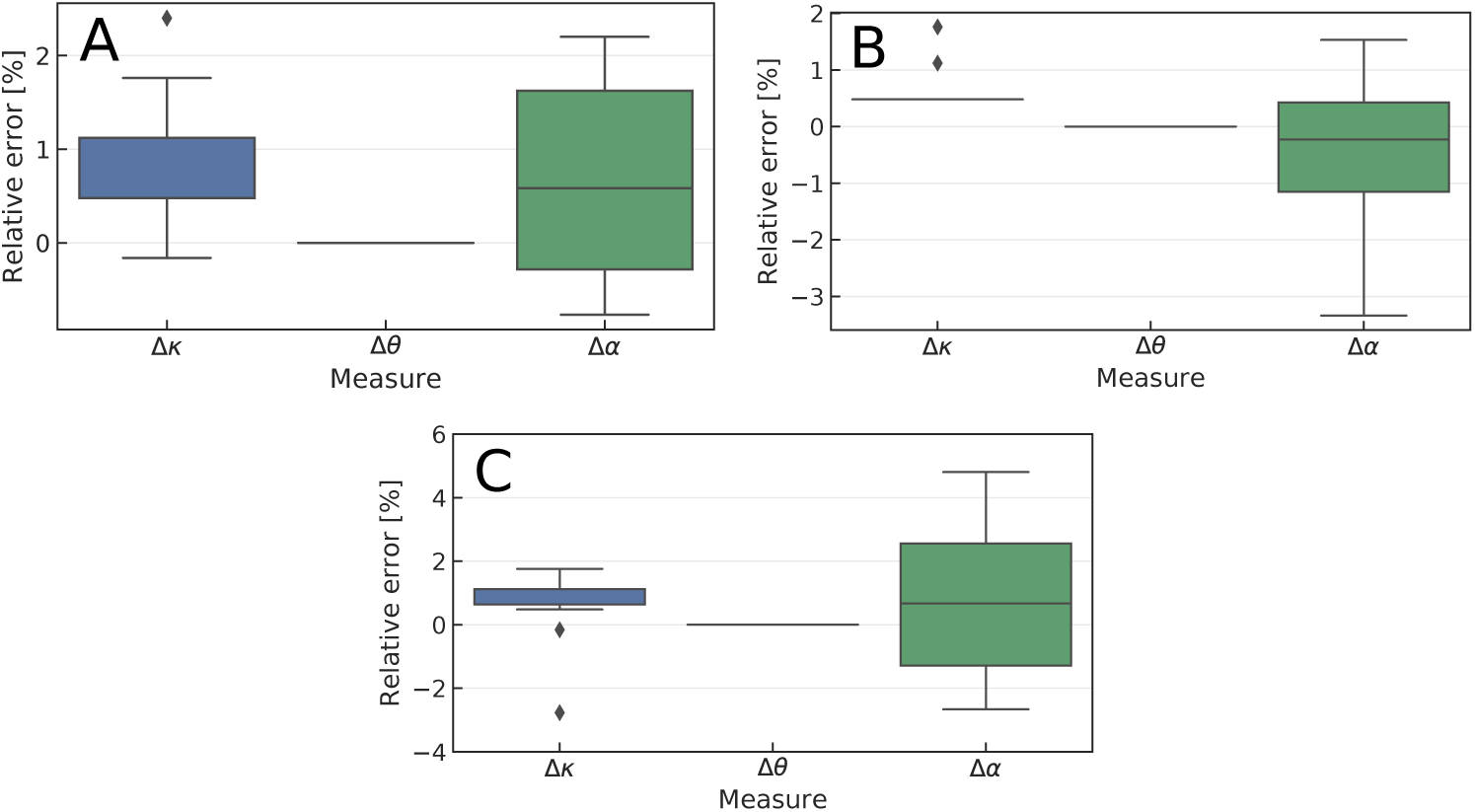
Relative errors of the transition to transversion ratio Δ k the heterozygosity Δ *θ*, and the shape parameter of the Γ distributed mutation rate heterogeneity Δ*α* The true shape parameter is *α* = 0.3 (A), *α* = 1.0 (B), and *α* = 5.0 (C), respectively.

The heterozygosity *θ* determines the level of polymorphism present in the species. We tested the accuracy of PoMo with increasing heterozygosity because for high *θ*s the model assumption of having boundary mutations only is violated. We observe low errors in branch score distance for heterozygosity values up to *θ*= 0.1 in analyses of species trees with twelve species with ten individuals each and a tree height of 3*N_e_* (Fig. 3).

**Figure 3:**
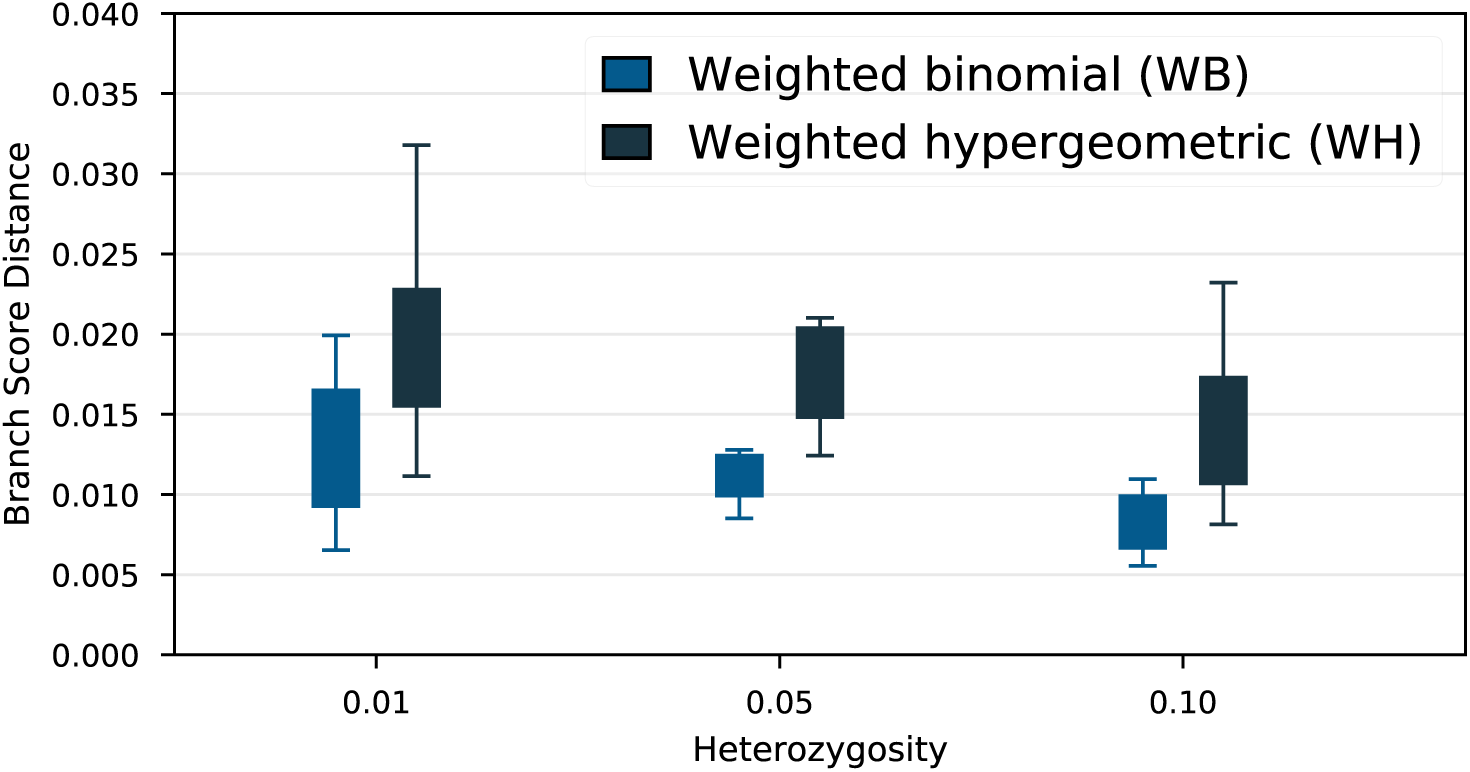
Branch score distance of weighted binomial and weighted hypergeometric sampling for Yule trees of height 3*N_e_* with twelve species and ten individuals each. Heterozygosity varies between 0.01 and 0.1.

For the case of uneven sampling of individuals of the populations, we tested two different strategies to account for the missing data in the poorly sampled populations. We found that weighted binomial sampling outperforms weighted hypergeometric sampling (see Materials and Methods and Supplement S3 for details on these strategies).

### ModelFinder and bootstrapping

Model selection and bootstrapping were tested on data of 12 great ape species (see Material and Methods). For PoMo, the GTR Tavaré (1986) mutation model with Γ rate heterogeneity was determined to be the best fitting model. The inferred Γ shape parameter is 1.26. The estimated phylogeny (Fig. 4) confirms previous results Schrempf *et al.* (2016). UFBoot2 and SH-aLRT both report 100 percent support for all branches.

**Figure 4:**
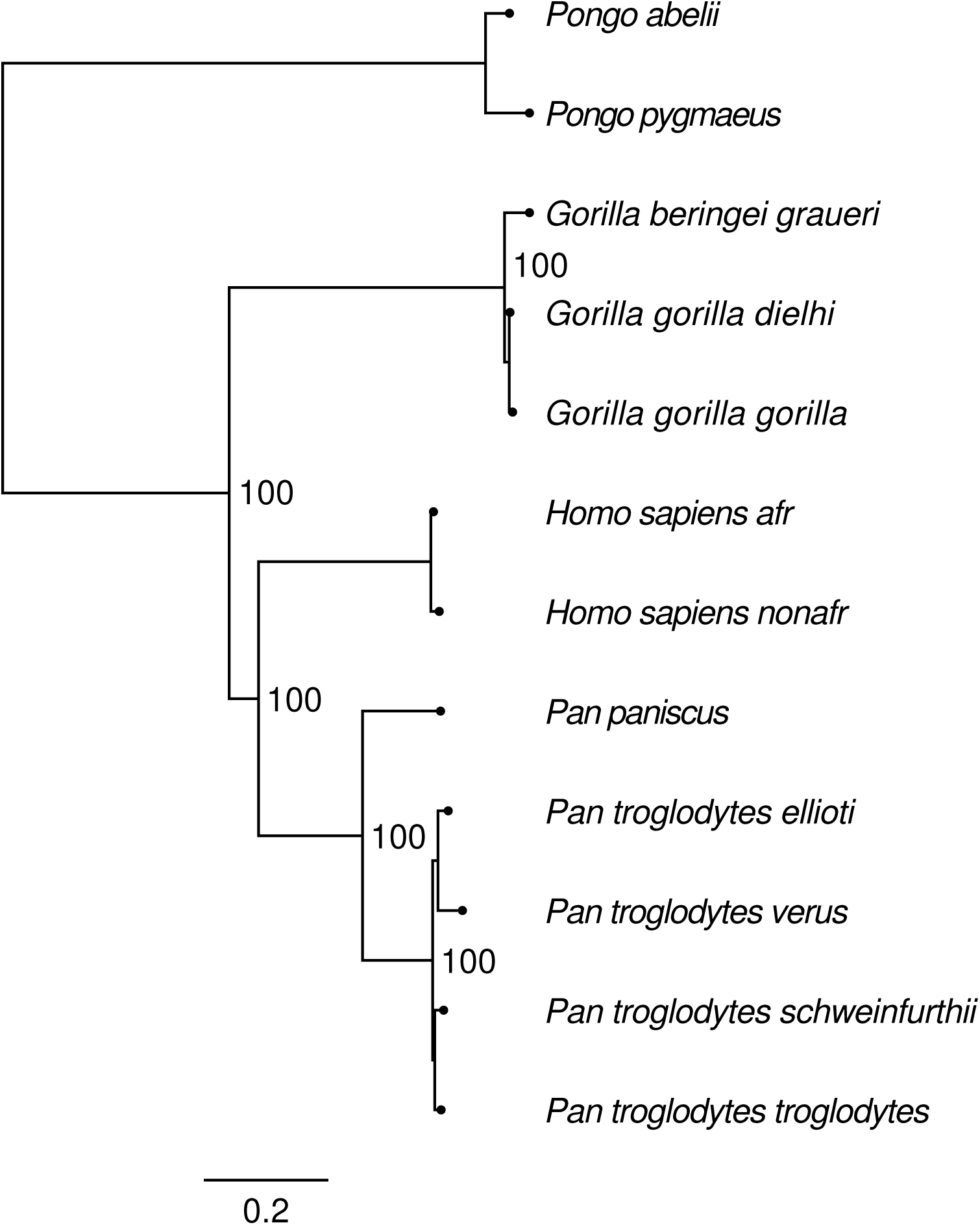
Phylogeny inferred from primate data Prado-Martinez *et al.* (2013). Both, UFBoot2 Hoang *et al.* (2018) and SH-aLRT Guindon *et al.* (2010) branch support tests evaluated to hundred percent support values.

## 4 Discussion

The analysis of data from 100 species underlines that IQ-TREE-PoMo is fast and scales well with increasing number of species. The chosen species tree exhibits significant amount of ILS, and hence, it is expected that our method which accounts for polymorphisms has higher accuracy than concatenation. In contrast to the concatenation method, which exhibits again statistical inconsistency (Fig. 1) as previously reported by Degnan and colleagues (Degnan, 2013; Degnan and Rosenberg, 2009), the branch score distance of IQ-TREE-PoMo continuously decreases when more data is analyzed (Fig. 1).

Further, it is now possible to infer parameters of more advanced sequence evolution models with IQ-TREE-PoMo. When modeling mutation rate heterogeneity, the shape parameter *α* of the Γ distribution is recovered with relative errors below two percent for 0.3 ≤ *α* ≤5.0. The inferred value of *α* = 1.26 for the great ape data set lies well in this range. We observe a slight overestimation of the shape parameter, when a significant amount of sites evolves extremely slowly and is nearly invariant (*α* ≤ 0.1, Fig. S2). Similarly and as expected, when the mutation rate distribution is homogeneous (*α*≥ 10.0), the variability of estimated shape parameters is higher (Fig. S4).

We decided not to implement the invariant sites model. The reason is that polymorphisms are the consequence of strictly positive mutation rates. When mutation rates are zero, all states but the boundary states have a stationary frequency of zero. This leads to problems because (1) observed polymorphic sites have zero likelihood and cannot be assigned to any model state and (2) numerical problems are encountered when calculating the transition probabilities. Although a separate model could be used to calculate the likelihood conditioned on the mutation rates being zero, we strongly believe that, if the data exhibits nearly invariant sites, those are sufficiently covered by Γ distributed mutation rate heterogeneity with low shape parameters.

As Mallo (2017) reported increased errors of the estimates of PoMo for high heterozygosity values up to *θ*= 0.05, we tested the robustness of our approach in this respect. We find that the accuracy of IQ-TREE-PoMo with respect to BSD increases with higher heterozygosity up to a value of *θ* = 0.1, which is well above the observed value in primates (*θ*~ 0.001, Prado-Martinez *et al.*, 2013) and most other organisms (except, Lynch *et al.*, 2016).

We recommend to use weighted binomial sampling instead of weighted hypergeometric sampling. Although weighted hypergeometric sampling has the advantage of retaining the heterozygosity between the observed data and the model, questions arise when the number of samples is lower than the number of PoMo frequency bins. In practice, the latter is often the cause of missing data. Correspondingly, IQ-TREE-PoMo uses weighted binomial sampling with *N* = 9 as default, a setting that has repeatedly been observed to be the most accurate and stable one Schrempf *et al.* (2016).

As more advanced mutation models are now available, the choice of the statistically most adequate mutation model is of major importance. ModelFinder does not require user input and is fast. The likelihood improvement when accounting for mutation rate heterogeneity is also tested for and, consequently, the most appropriate mutation rate heterogeneity model is reported and automatically used.

For the great ape data set, the best-fit model for PoMo coincides with results from concatenation methods and from the phylogenetic analysis in the original paper Prado-Martinez *et al.* (2013). In the latter case, not only the model choice coincides, but also the topology, which is a further confirmation of the validity of PoMo.

Additionally, assessment of branch support values is now possible with normal bootstrap, UFBoot2 and likelihood ration based tests (SH-aLRT). Branch support values for the primate data are high because 2.8 million sites are considered. This is not surprising and emphasizes that the results are robust. Especially for smaller data sets, this feature will be highly useful.

Overall, we provide important extensions to further establish the use of polymorphism-aware models in phylogenetics. In the future, we would like to implement probability distribution free rate category models Kalyaanamoorthy *et al.* (2017). We have repeatedly discussed allowing non-reversible mutation models to be used with PoMo since IQ-TREE offers a whole set of non-reversible mutation models consistent with heterogeneous mutation rates (Lie-Markov models, Woodhams *et al.*, 2015). IQ-TREE does not only provide an extensive set of mutation models or allows usage of new model selection methods such as ModelFinder, but also comes with other benefits such as the ability to resume analyses (checkpointing), or carefully designed parallelization techniques. In conclusion, PoMo has evolved to be a mature, well-tested and flexible method to perform phylogenetic inference with population data.

## 5 Materials and Methods

### PoMo

A detailed description of PoMo and the discrete multivariate boundary mutation model can be found in Schrempf *et al.* (2016) and Schrempf and Hobolth (2017), respectively. PoMo is a time-continuous Markov process modeling sequence evolution along a species tree. Sites are assumed to be independent (composite likelihood, free recombination). At each site, not only evolution of a single character *a*∈ *A*(*e*.g., *A*= {*A, C, G, T* | *A* |= 4) of the reference genome is considered but rather the evolution of the collective characters of a population of genomes.

Actual populations are big in size and direct treatment is not feasible. For neutral evolution, effective population size is confined with mutation rates. It is possible to scale down the effective population size to a small value while scaling up mutation rates such that the overall dynamics remain unchanged. We take advantage of this property by choosing a rather small number of collected characters *N* of the multivariate boundary mutation model. Consequently, the parameter *N* should be interpreted as a discretization parameter (not to be confused with effective population size) describing the number of bins that allele frequencies can fall into.

The rates of frequency shifts, are determined using the time-continuous Moran process Moran (1958). PoMo assumes that drift removes variation fast, and mutations are disallowed when more than one allele is present in the population. Hence, the population can either be monomorphic for an allele *a*∈ ***A***or polymorphic for two alleles *a, b* ∈ ***A*** with counts *i* and (*N-i*), respectively. The monomorphic and polymorphic states of the multivariate boundary mutation model are denoted by {a} and {*ia* | (*N-i*)*b}*, *a* #*b*, respectively. Disallowing mutations when the population is polymorphic is a good approximation as long as the heterozygosity is below a value of 0.1, a requirement that is readily satisfied in most cases.

The transition rate matrix

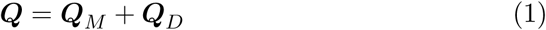

of dimension 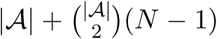 is composed by mutation (M) and genetic drift (D). The only off-diagonal, non-zero entries of ***Q****_M_* are mutations away from monomorphic states

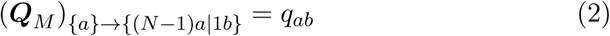

where the *q_ab_* are mutation rates defined by an underlying mutation model such as the HKY mutation model Hasegawa *et al.* (1985), or the GTR mutation model Tavaré (1986). Drift rates are non-zero for neighboring states only (1 ≤*i* ≤N-1)

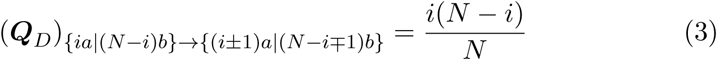

Diagonal elements of ***Q****_M_* and ***Q****_D_* are set such that all row sums are zero.

### Big data

For the simulation of large data sets, we employed a pipeline that follows Schrempf *et al.* (2016) First, ten species trees with 100 leaves were randomly generated under the Yule birth model Yule (1925). Each of the ten species trees is referred to as one of ten replicates. The height of the species trees measured in number of generations was 6 times the effective population size which is assumed constant. The Yule birth rate was set such that the expected number of species for the given height is 100. Second, for each replicate, 1000 gene trees were simulated under the multispecies coalescent model. SimPhy Mallo *et al.* (2016) was used for these steps. Finally, for each gene tree, DNA sequences with 1000 sites were generated with Seq-Gen Rambaut and Grass (1997) under the HKY mutation model Hasegawa *et al.* (1985). The transition to transversion ratio was *κ*= 6.25, the stationary nucleotide frequencies were *π_A_*= 0.3, *π_C_* = 0.2, *π_G_*= 0.2, and *π_T_* = 0.3. The simulated sequences had a heterozygosity of 0.005 which is approximately four times the value observed in primates (e.g., Prado-Martinez *et al.*, 2013).

Estimation of the original species trees with 100 leaves was performed with IQ-TREE using either PoMo or a standard concatenation method. The HKY model was used for both methods. The discretization parameter of the multivariate boundary mutation model was *N* = 10 and weighted binomial sampling was used. Command lines for simulation and analysis are in Section S1. The accuracy of the inferences was measured by comparing the estimated to the original species trees. Trees were normalized to a tree height of 1.0. The branch score distance (BSD, Kuhner and Felsenstein, 1994) and the Robinson-Foulds distance Robinson and Foulds (1981) were used.

### High heterozygosity

For the assessment of the accuracy of tree inference for different heterozygosity values (Fig. 3) the same simulation pipeline was used. In contrast to above, the height of the species trees was 3 times the effective population size, the number of taxa was twelve, and the heterozygosity values was 0.01, 0.05, and 0.1, respectively.

### Simulator

Although the above pipeline corresponds to the way how methods based on the multispecies coalescent model perform inference, estimates of PoMo are accurate (Fig. 3 and Schrempf *et al.*, 2016). When considering mutation rate heterogeneity, we have to keep in mind that the mutation rates of the multivariate boundary mutation model are not independent of the effective population size. A mutational event {a}→ {(*N -* 1)*a*p 1*b* }, *a, b* ∈ ***A***,*a* ≠ *b*, corresponds to a mutation in the real population with subsequent frequency shift to a value of 1*/N*. Of course, the probability of such an event depends on the effective population size of the real population. As a result, higher mutation rates in the multivariate boundary mutation model, associated with a higher probability to be in a polymorphic state when species split, lead to a higher probability of ILS.

In contrast, the probability of ILS derived by the multispecies coalescent model does not explicitly depend on the mutation rate. For example, a rooted, three-taxon tree has a probability of ILS proportional to *e*^−*c*Δt/*N_e_*^ (e.g., Nei, 1987), where *c* is a constant, Δ*t* is the branch length of the internal branch measured in number of generations, and *N_e_* is the effective population size. These considerations urged us to develop a multivariate boundary mutation model simulator for direct assessment of the accuracy of mutation rate heterogeneity inference.

### Advanced models

Heterogeneity in evolutionary rates across sites may be modeled with a parametric distribution such as the Γ distribution Yang (1994). For the multivariate boundary mutation model, complications arise because the transition rate matrix contains contributions from mutations ***Q****_M_* as well as frequency shifts due to random genetic drift ***Q****_D_*. A general scaling of the transition rate matrix, like it is usually done when using Γ rate heterogeneity with substitution models, would erroneously (de-)accelerate both processes. This difficulty was overcome by employing the concept of mixture models.

In brief, *K* mutation rate categories with rate modifiers *r_k_* and corresponding probabilities 1*/K* of a site belonging to a category are defined. The total likelihood of a mutation model accounting for mutation rate heterogeneity *M_h_* and species tree *T* given data *D* is

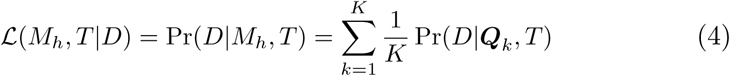

where

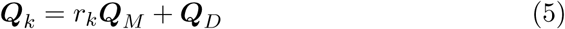

Hereby, the mutation rates modifiers *r_k_* are the means of the mutation rate categories. The latter is similar to the classical treatment using Γ rate heterogeneity, however, this setup allows modeling of mutation rate heterogeneity with any free mutation rate categories. We chose to use a parametrized Γ distribution because it is most used. Naturally, the usual efficiency of Γ rate heterogeneous models is not retained, because the transition matrices for different mutation rate categories are intrinsically different. That is, an analysis with *k* categories requires eigendecomposition of *k* transition rate matrices.

The accuracy in estimating the shape parameter of the Γ distributed mutation rate heterogeneity was assayed using bmm-simulate. Species trees were randomly sampled from a Yule process. The tree height measured in average number of substitutions per site was *h* = 0.01. The speciation rateλ was set such that *n* = 12 species are present on average. That is (e.g., Kendall, 1949)

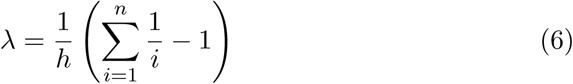

For each sampled species tree, sequences were simulated under the discrete multivariate boundary mutation model. The discretization parameter *N* was set to 10, and the heterozygosity to 0.0025. Similar to the big data analysis above, the HKY mutation model was used with *κ* = 6.25 and stationary nucleotide frequencies *π_A_* = 0.3, *π_C_*= 0.2, *π_G_* = 0.2, and *π_T_* = 0.3. Four Γ mutation rate categories were used and one million sites were simulated. Ten replicate analyses with different, randomly sampled species trees were performed. The shape parameter *α* of the Γ distributed mutation rate heterogeneity was set to 0.1, 0.3, 0.5, 1.0, 5.0, and 10.0. Command lines for simulation and analysis with rate heterogeneity are in Section S2.

### Sampling

Interpretation of data is mostly predetermined when using phylogenetic substitution models, because observed states can directly be mapped to internal states of the used substitution model. Special handling is required when encountering an unknown site. Then, the likelihood of all states is set to 1.0, and all model states have the same probability of leading to the observed data.

Equivalently, we would like to assign the likelihoods of all multivariate boundary model states at a specific site in the alignment and leaf of the tree. If the number of samples coincides with the discretization parameter *N*, and if not more than two alleles are present, the observed data is a multivariate boundary mutation model state. Like above with substitution models, the likelihood of this state can be set to 1.0, and the likelihood of all other states to 0.0.

In practice, we often do not have total control over the number of individuals sampled. For example, sequences are retrieved from existing databases, or the high sequencing costs set constraints on the number of individuals. We therefore designed various strategies to assign the likelihoods of the multivariate boundary mutation model states Schrempf *et al.* (2016) if the number of samples does not match *N*. The easiest one is to randomly draw *N* alleles with replacement from the data. We refer to this methods as *sampled*. We can also assign likelihoods in an inverse manner, similar to handling unknown sites with substitution models. A specific multivariate boundary mutation model state is assumed at the considered leaf and the probability of observing the data given this state is calculated. We can sample with or without replacement and calculate this probability for all multivariate boundary mutation model states. We have termed these ways of assigning likelihoods *weighted binomial* and *weighted hypergeometric* sampling. A detailed description is given in Section S3.

### Bootstrapping using real data

We revisited the great ape data set of six species subdivided into 12 populations with up to 23 individuals each Prado-Martinez *et al.* (2013). The exome-wide data set includes roughly 2.8 million, 4-fold degenerate sites. Data preparation is described in De Maio *et al.* (2015), the counts file is available on https://github.com/pomo-dev/data.

For the analysis we used ModelFinder and assessed branch support with 1000 bootstraps. We tested estimations with and without Γ mutation rate heterogeneity with four discrete categories combined with UFBoot2 and SH-aLRT. The discretization parameter was *N* = 9. The used command line was

`iqtree -s hg18-all.cf -alrt 1000 -bb 1000`

## Supporting information

## 6 Acknowledgments

This work was funded by the Vienna Science and Technology Fund (WWTF) through project MA16-061. DS was supported by the Austrian Science Fund [FWF-P24551, I-2805-B29] and received funding from the European Research Council under the European Union’s Horizon 2020 research and innovation programme under grant agreement no. 741774. The computational results presented have been achieved in part using the Vienna Scientific Cluster (VSC) and the St Andrews Bioinformatics Unit (StABU) which is funded by a Wellcome Trust ISSF award (grant 105621/Z/14/Z).

