## Supplementary material for "Polymorphism-aware species trees with advanced mutation models, bootstrap and rate heterogeneity"

November 26, 2018

### Contents

**S1 Big data** **2**

**S2 Simulator and rate heterogeneity** **4**

**S3 Sampling strategy** **6**

### S1 Big data

Command lines used for simulation and analysis pipeline.

1. Generate gene trees with **SimPhy**.

```
simphy -sb f:0.0006978962529399367 -rs 20 -rl f:1 \
  -rg 1000 -sp f:1000 -sg f:1 -sl f:100 -st f:6000 \
  -si f:10 -su f:0.001 \
  -o Yule100Tree_6NeHeight_10Indivs_1000Genes
```

2. Generated gene trees need to be extracted into separate files (e.g. with Python, not shown here).

3. Generate multi-sequence alignment for each gene tree using **SeqGen**.

```
seq-gen -mHKY -f0.3,0.2,0.2,0.3 -t3.0 -l1000 \
  -n1 -on -s0.0025
  < Yule100Tree_6NeHeight_10Indivs_Tree_Gene1 \
  > Yule100Tree_6NeHeight_10Indivs_MSA_Gene1
```

4a. Concatenation of all genes (e.g. with Python, not shown here). Subsequent analysis with concatenation method.

```
iqtree -m HKY \
  -s Yule100Tree_6NeHeight_10Indivs_MSA_AllGenes
```

4b. Preparation of counts files (e.g. with `cflib` from `https://github.com/pomo-dev/cflib`, not shown here). Subsequent analysis with IQ-TREE PoMo.

```
iqtree -m HKY+P+N10+WB \
  -s Yule100Tree_6NeHeight_10Indivis_CountsFile_AllGenes
```

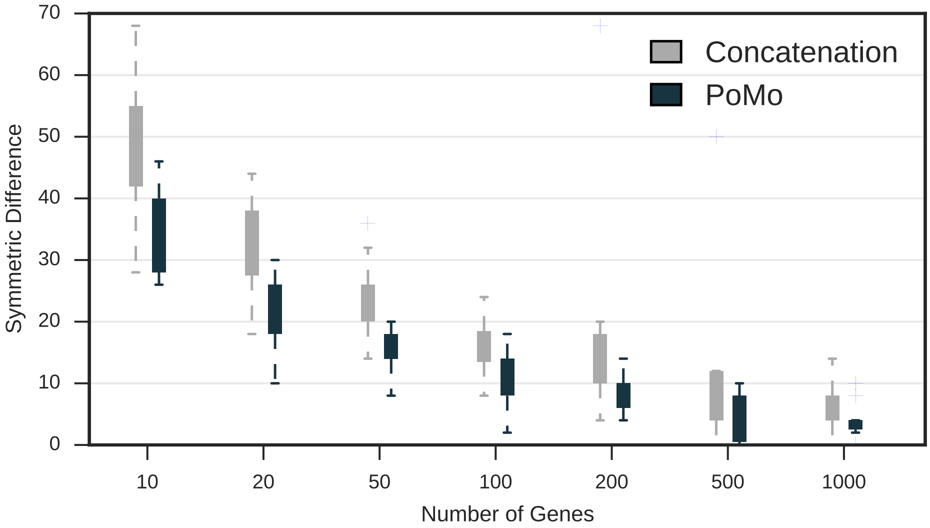


**Figure S1:** Symmetric difference or Robinson-Foulds metric (Robinson and Foulds, 1981) of concatenation approach and IQ-TREE PoMo with $N = 10$ and weighted binomial sampling for the Yule trees with 100 species and 10 samples each. The tree height is $6N_e$. The HKY model was used for both inference methods. The heterozygosity is $\theta_W = 0.005$ per site. Each gene spans 1000 sites. The error bars are standard deviations of ten replicate analyses.

#### S2 Simulator and rate heterogeneity

Command lines used for simulation and analysis pipeline.

1. For example, generate counts files with rate heterozygosity with Γ shape parameter $\alpha = 1.0$.

```
bmm-simulate -N 10 --heterozygosity 0.0025 \
  --tree-type Yule --tree-height 0.01 \
  --tree-yule-rate 210.3210678210678 \
  --gamma-shape 1.0 --gamma-ncat 4 \
  --nsites 1000000 -o data.cf
```

2. Analysis with HKY model, rate heterogeneity, $N = 10$, and weighted hypergeometric sampling.

```
iqtree -s data.cf -m HKY+P+G4+N10+WH
```

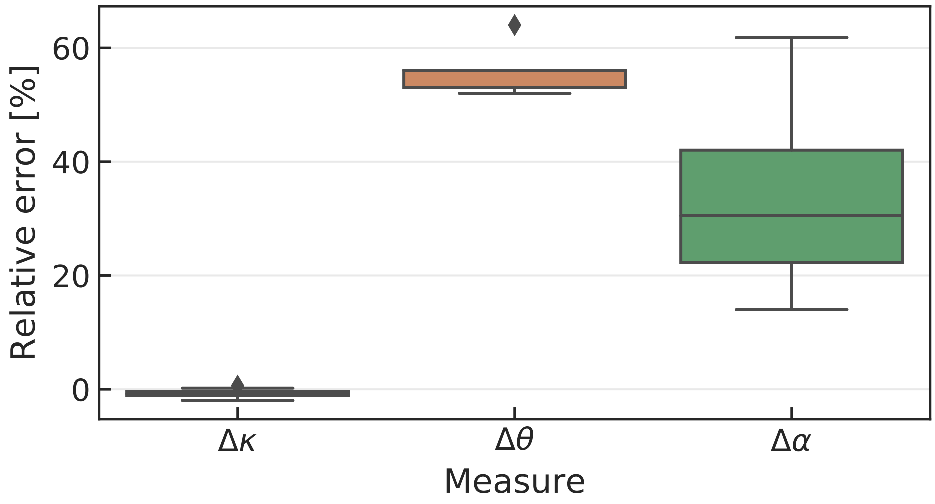


**Figure S2:** Relative errors of the transition to transversion ratio $\Delta\kappa$, the heterozygosity $\Delta\theta$ and the shape parameter of the Γ distributed mutation rate heterogeneity $\Delta\alpha$ for a Yule tree of height 0.01 average number of substitutions per site and an average number of twelve species with ten samples each. One million sites where analyzed, the true heterozygosity and shape parameter are $\theta = 0.0025$ and $\alpha = 0.1$, respectively.

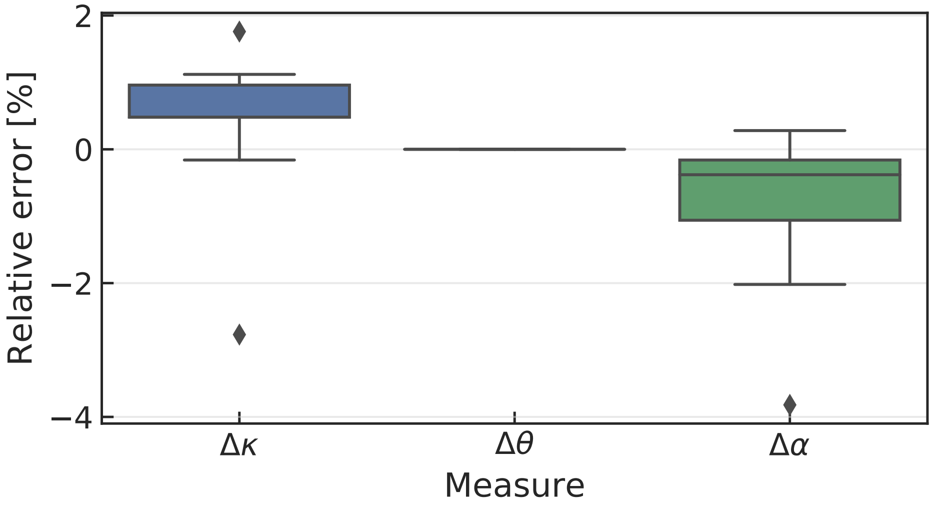


**Figure S3:** Relative errors of the transition to transversion ratio $\Delta\kappa$, the heterozygosity $\Delta\theta$ and the shape parameter of the $\Gamma$ distributed mutation rate heterogeneity $\Delta\alpha$ for a Yule tree of height 0.01 average number of substitutions per site and an average number of twelve species with ten samples each. One million sites where analyzed, the true heterozygosity and shape parameter are $\theta = 0.0025$ and $\alpha = 0.5$, respectively.

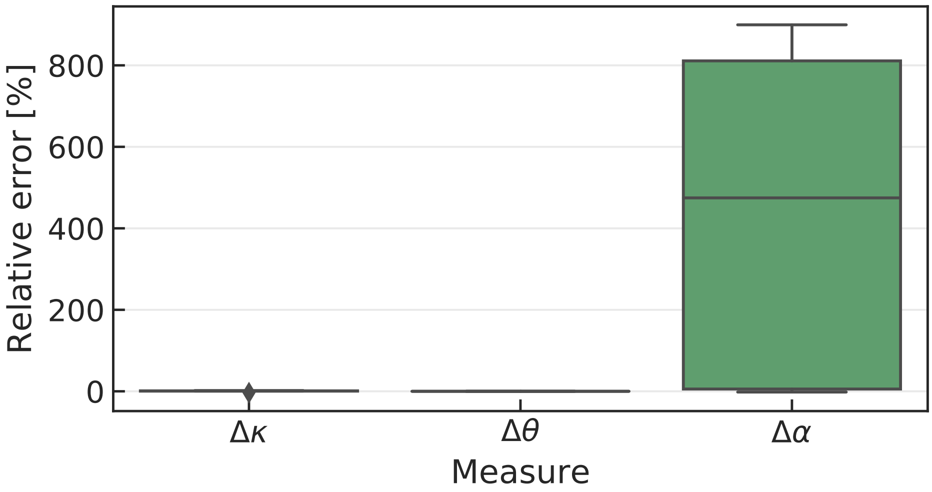


**Figure S4:** Relative errors of the transition to transversion ratio $\Delta\kappa$, the heterozygosity $\Delta\theta$ and the shape parameter of the $\Gamma$ distributed mutation rate heterogeneity $\Delta\alpha$ for a Yule tree of height 0.01 average number of substitutions per site and an average number of twelve species with ten samples each. One million sites where analyzed, the true heterozygosity and shape parameter are $\theta = 0.0025$ and $\alpha = 10$, respectively.

#### S3 Sampling strategy

Interpretation of data is mostly predetermined when using phylogenetic substitution models, because observed states can directly be mapped to internal states of the used substitution model. Let the state space of the considered substitution model be nucleotide bases $\mathcal{A} = \{A, C, G, T\}$. If, at a specific site in the alignment and leaf of the tree, we observe an $A$, the likelihood vector will be set to ($a \in \mathcal{A}$)

$$\mathcal{L}(a|A) = \begin{cases} 1 & \text{if } a = A, \\ 0 & \text{otherwise.} \end{cases} \tag{S1}$$

Special handling is required when encountering an unknown site (usually denoted by $N$, not to be confused with the discretization parameter of the multivariate boundary mutation model). Then, likelihoods are assigned in an inverse manner. That is, the probability of a specific state leading to the observed data is considered. In this example, the probability is set to 1, irrespective of the nucleotide. I.e., any state at the tip is equally likely

$$\mathcal{L}(a|N) = 1, \quad a \in \mathcal{A}. \tag{S2}$$

Equivalently, we would like to assign the likelihoods of all multivariate boundary mutation model states at a specific site in the alignment and leaf of the tree. If the number of samples coincides with the discretization parameter $N$, and if not more than two alleles are present, the observed data is a multivariate boundary mutation model state. Like above with substitution models, the likelihood of this state can be set to 1, and the likelihood of all other states to 0. Let us suppose that $N = 10$ and that we observe 4 individuals with an $A$ and 6 individuals with a $C$. Then, $x \in \mathcal{A}_{BM}$, where $\mathcal{A}_{BM}$ is the state space of the multivariate boundary mutation model

$$\mathcal{L}(x|\{4A|6C\}) = \begin{cases} 1 & \text{if } x = \{4A|6C\}, \\ 0 & \text{otherwise.} \end{cases} \tag{S3}$$

However, if the number of samples does not match the discretization parameter $N$, and this will often be the case, we have to think about other means of assigning the likelihoods of the multivariate boundary mutation model states Schrempf *et al.* (2016). The easiest one is to randomly draw $N$ alleles with replacement from the data (each allele has equal probability of being chosen; binomial sampling). We refer to this methods as *sampled*. Let us suppose that we observe that 13 individuals have an $A$ and 18 individuals have a $C$, i.e., $\{13A|18C\} \notin \mathcal{A}_{BM}$. So, we randomly pick $N$ alleles from the observed data. Let us suppose that $N = 10$ and that we pick 3 individuals with an $A$ and 7 individuals with a $C$. So ($x \in \mathcal{A}_{BM}$)

$$\mathcal{L}(x|\{3A|7C\} \text{ sampled from } \{13A|18C\}) = \begin{cases} 1 & \text{if } x = \{3A|7C\}, \\ 0 & \text{otherwise.} \end{cases} \tag{S4}$$

A more canonical way is to reverse reasoning, similar to handling unknown sites with substitution models. Let us fix a specific multivariate boundary mutation model state at the considered site and leaf and let us consider the probability of observing the data given this state. There are many ways of defining this probability. The simplest is, (1) assume that all alleles have equal probability of being drawn, and (2) sample with (binomial) or without (hypergeometric) replacement. We do this for all multivariate boundary mutation model states. We have termed these ways of assigning likelihoods *weighted binomial* or *weighted hypergeometric* sampling.

For example, let the observed data be the same as before $\{13A|18C\}$, and let us use weighted binomial sampling. Then, the likelihood of multivariate boundary mutation model state $\{1A|9C\}$ would be

$$\mathcal{L}(\{1A|9C\}|\{13A|18C\}) = \Pr(\{13A|18C\}|\{1A|9C\}) \tag{S5}$$

$$= \binom{31}{13}\left(\frac{1}{10}\right)^{13}\left(\frac{9}{10}\right)^{18}. \tag{S6}$$

In general, when observing $M$ alleles at the considered site and leaf ($0 \leq i \leq N$; $0 \leq j \leq M$; $a, b, c, d \in \mathcal{A}$)

$$\mathcal{L}(\{ia|(N-i)b\}|\{jc|(M-j)d\}) = \begin{cases} \binom{M}{j}\left(\frac{i}{N}\right)^{j}\left(\frac{N-i}{N}\right)^{M-j} & \text{if } a = c \text{ and } b = d, \\ 0 & \text{otherwise.} \end{cases} \tag{S7}$$
